# Pharmacologic Activation of Integrated Stress Response Kinases Inhibits Pathologic Mitochondrial Fragmentation

**DOI:** 10.1101/2024.06.10.598126

**Authors:** Kelsey R. Baron, Samantha Oviedo, Sophia Krasny, Mashiat Zaman, Rama Aldakhlallah, Prerona Bora, Prakhyat Mathur, Gerald Pfeffer, Michael J. Bollong, Timothy E. Shutt, Danielle A. Grotjahn, R. Luke Wiseman

## Abstract

Excessive mitochondrial fragmentation is associated with the pathologic mitochondrial dysfunction implicated in the pathogenesis of etiologically-diverse diseases, including many neurodegenerative disorders. The integrated stress response (ISR) – comprising the four eIF2α kinases PERK, GCN2, PKR, and HRI – is a prominent stress-responsive signaling pathway that regulates mitochondrial morphology and function in response to diverse types of pathologic insult. This suggests that pharmacologic activation of the ISR represents a potential strategy to mitigate pathologic mitochondrial fragmentation associated with human disease. Here, we show that pharmacologic activation of the ISR kinases HRI or GCN2 promotes adaptive mitochondrial elongation and prevents mitochondrial fragmentation induced by the calcium ionophore ionomycin. Further, we show that pharmacologic activation of the ISR reduces mitochondrial fragmentation and restores basal mitochondrial morphology in patient fibroblasts expressing the pathogenic D414V variant of the pro-fusion mitochondrial GTPase MFN2 associated with neurological dysfunctions including ataxia, optic atrophy, and sensorineural hearing loss. These results identify pharmacologic activation of ISR kinases as a potential strategy to prevent pathologic mitochondrial fragmentation induced by disease-relevant chemical and genetic insults, further motivating the pursuit of highly selective ISR kinase-activating compounds as a therapeutic strategy to mitigate mitochondrial dysfunction implicated in diverse human diseases.

## INTRODUCTION

The integrated stress response (ISR) comprises four stress-regulated kinases – PERK, PKR, GCN2, and HRI – that selectively phosphorylate eIF2α in response to diverse pathologic insults.^1,2^ The ISR has recently emerged as a prominent stress-responsive signaling pathway activated by different types of mitochondrial stress.^3–5^ Mitochondrial uncoupling or inhibition of ATP synthase activates the ISR through a mechanism involving the cytosolic accumulation and oligomerization of the mitochondrial protein DELE1, which binds to and activates the ISR kinase HRI.^6–10^ Complex I inhibition can also activate the ISR downstream of the ISR kinase GCN2.^11^ Further, CRISPRi-depletion of mitochondrial proteostasis factors preferentially activates the ISR over other stress-responsive signaling pathways.^12^ These results identify the ISR as an important stress-responsive signaling pathway activated in response to many different mitochondrial insults.

Consistent with its activation by mitochondrial stress, the ISR regulates many aspects of mitochondrial biology. ISR kinases are activated in response to stress through a conserved mechanism involving dimerization and autophosphorylation.^1,2^ Once activated, these kinases phosphorylate eIF2α, resulting in both a transient attenuation of new protein synthesis and the activation of stress-responsive transcription factors such as ATF4.^1,2^ The ISR promotes adaptive remodeling of mitochondrial pathways through both transcriptional and translational signaling. For example, the activation of ATF4 downstream of phosphorylated eIF2α transcriptionally regulates the expression of mitochondrial chaperones (e.g., the mitochondrial HSP70 *HSPA9*) and proteases (e.g., the AAA+ protease *LON*), increasing mitochondrial proteostasis capacity during stress.^13,14^ ISR-dependent translational attenuation also enhances mitochondrial proteostasis through the selective degradation of the core import subunit TIM17A, which slows mitochondrial protein import and reduces the load of newly-synthesized proteins entering mitochondria during stress.^15^ Apart from proteostasis, transcriptional and translational signaling induced by the activation of ISR kinases also regulates many other mitochondrial functions, including phospholipid synthesis, cristae organization, and electron transport chain activity.^16–20^

Intriguingly, the morphology of mitochondria is also regulated by ISR signaling. Mitochondrial morphology is dictated by the relative activities of pro-fission and pro-fusion GTPases localized to the outer and inner mitochondrial membranes (OMM and IMM, respectively).^21,22^ These include the pro-fusion GTPases MFN1 and MFN2 and the pro-fission GTPase DRP1, all localized to the OMM. Previous results showed that ER stress promotes adaptive mitochondrial elongation downstream of the PERK arm of the ISR through a mechanism involving the accumulation of the phospholipid phosphatidic acid on the OMM where it inhibits the pro-fission GTPase DRP1.^19,20,23^ This PERK-dependent increase of mitochondrial elongation functions to protect mitochondria during ER stress through multiple mechanisms, including enhanced regulation of respiratory chain activity, suppression of mitochondrial fragmentation, and reductions in the turnover of mitochondria by mitophagy.^19,20,23^ Apart from PERK, pharmacologic activation of the alternative ISR kinase GCN2 also promotes adaptive mitochondrial elongation.^23^ This indicates that mitochondrial elongation may be a protective mechanism that can be pharmacologically accessed by activating multiple different ISR kinases.

Excessive mitochondrial fragmentation is a pathologic hallmark of numerous human diseases, including several neurodegenerative disorders.^21,24–27^ Fragmented mitochondria are often associated with mitochondrial dysfunctions including impaired respiratory chain activity and dysregulation of mitophagy.^21,24–29^ Thus, pathologic increases in mitochondrial fragmentation are linked to many mitochondrial dysfunctions implicated in human disease. Consistent with this observation, interventions that prevent mitochondrial fragmentation have been shown to correct pathologic mitochondrial dysfunction in models of many different diseases.^30,31^ Notably, genetic or pharmacologic inhibition of the pro-fission GTPase DRP1 blocks pathologic mitochondrial fragmentation and subsequent mitochondrial dysfunction in cellular and in vivo models of diverse neurodegenerative diseases, including Charcot-Marie-Tooth Type 2A and Type 2B, spastic paraplegia, optic atrophy, Huntington’s disease, Amyotrophic Lateral Sclerosis, and Alzheimer’s disease.^32–43^

The ability of ISR activation to promote mitochondrial elongation suggests that pharmacologic activation of ISR kinases could mitigate the pathologic mitochondrial fragmentation implicated in etiologically diverse human diseases. To test this idea, we probed the potential for pharmacologic activation of different ISR kinases to reduce mitochondrial fragmentation induced by disease-relevant chemical and genetic insults. Previous work identified the small molecule halofuginone as a potent activator of the ISR kinase GCN2 that promotes ISR-dependent, adaptive mitochondrial elongation.^23,44^ However, few other compounds were available that selectively activated other ISR kinases and were suitable for probing mitochondrial adaptation induced by ISR kinase activation.^23,45^ Here, we used a small molecule screening platform to identify two nucleoside mimetic compounds, 0357 and 3610, that preferentially activate the ISR downstream of the ISR kinase HRI. Using these compounds, we demonstrate that pharmacologic HRI activators also promote adaptive, ISR-dependent mitochondrial elongation. We go on to show that pharmacologic activation of the ISR with these compounds prevents DRP1-mediated mitochondrial fragmentation induced by the calcium ionophore ionomycin.^46,47^ Further, we show that compound-dependent activation of the ISR reduces the population of fragmented mitochondria and rescues basal mitochondrial network morphology in patient fibroblasts expressing the D414V variant of the pro-fusion GTPase MFN2 associated with a complex clinical phenotype including ataxia, optic atrophy and sensorineural hearing loss.^48^ These results demonstrate the potential for pharmacologic activation of the ISR to prevent pathologic mitochondrial fragmentation in disease-relevant models. Moreover, our work further motivates the continued development of highly selective activators of ISR kinases as a potential therapeutic strategy to mitigate the pathologic mitochondrial dysfunction implicated in etiologically diverse human diseases.

## RESULTS

### The nucleoside mimetic compounds 0357 and 3610 preferentially activate the ISR downstream of HRI

The small molecule halofuginone activates the ISR kinase GCN2 and induces ISR-dependent mitochondrial elongation.^23,44^ However, few other compounds are available that selectively activate other ISR kinases through mechanisms that allow for ISR-dependent mitochondrial remodeling.^23^ For example, BtdCPU activates the ISR kinase HRI through a mechanism involving mitochondrial uncoupling, precluding its use for probing ISR-dependent protection of mitochondria.^23^ To address this limitation and define the potential for pharmacologic activation of other ISR kinases to promote adaptive mitochondrial elongation, we established and implemented a screening platform to identify compounds that activated ISR signaling downstream of alternative ISR kinases (**Fig. 1A**). In this screen, we used the ATF4-FLuc translational reporter of the ISR (**Fig. S1A**).^10^ We confirmed that ISR activating stressors including the ER stressor thapsigargin (Tg) and the ATP synthase inhibitor oligomycin A (OA) robustly activated this reporter (**Fig. S1B**). We then used this reporter to screen the ∼3k nucleoside mimetic analog compound library (10 µM) and monitored ATF4-FLuc activity 8 h after treatment. Our primary screen identified 34 hit compounds that activated the ATF4-FLuc reporter with a robust Z-score >3-fold. We then removed highly reactive compounds and pan-assay interference compounds (PAINS), which reduced the number of hits to 9 (**Fig. 1B**). These compounds were re-purchased and then tested in dose response for ATF4-FLuc activation (**Fig. S1C**). This identified compounds 0357 and 3610 as the compounds that most efficaciously activated the ATF4-FLuc reporter, albeit with low potency (EC_50_ > 10 µM). We confirmed that co-treatment with the highly selective ISR inhibitor ISRIB^49,50^ blocked ATF4-FLuc activation induced by these compounds, confirming this activation can be attributed to the ISR (**Fig. 1C,D**). Further, we used qPCR to show that treatment with 0357 or 3610 increased expression of the ISR target genes *ASNS* and *CHAC1* in HEK293 and MEF cells (**Fig. S1D,E**).^51,52^ Importantly, these compounds did not activate luciferase reporters of other stress-responsive signaling pathway such as the unfolded protein response (UPR; XBP1-RLuc)^53,54^, the heat shock response (HSR; HSE-FLuc)^55^, or the oxidative stress response (OSR; ARE-FLuc)^56^ (**Fig. 1E,F**). Further, treatment with these compounds did not induce expression of the UPR target gene *BiP*, the HSR target gene *HSPA1A*, or the OSR target gene *NQO1* in HEK293 cells (**Fig. S1F**). Finally, treatment with 0357 or 3610 did not significantly reduce viability of HEK293 cells (**Fig. S1G**). These results indicate compounds 0357 and 3610 preferentially activate the ISR, as compared to other stress responsive signaling pathways.

**Figure 1.**
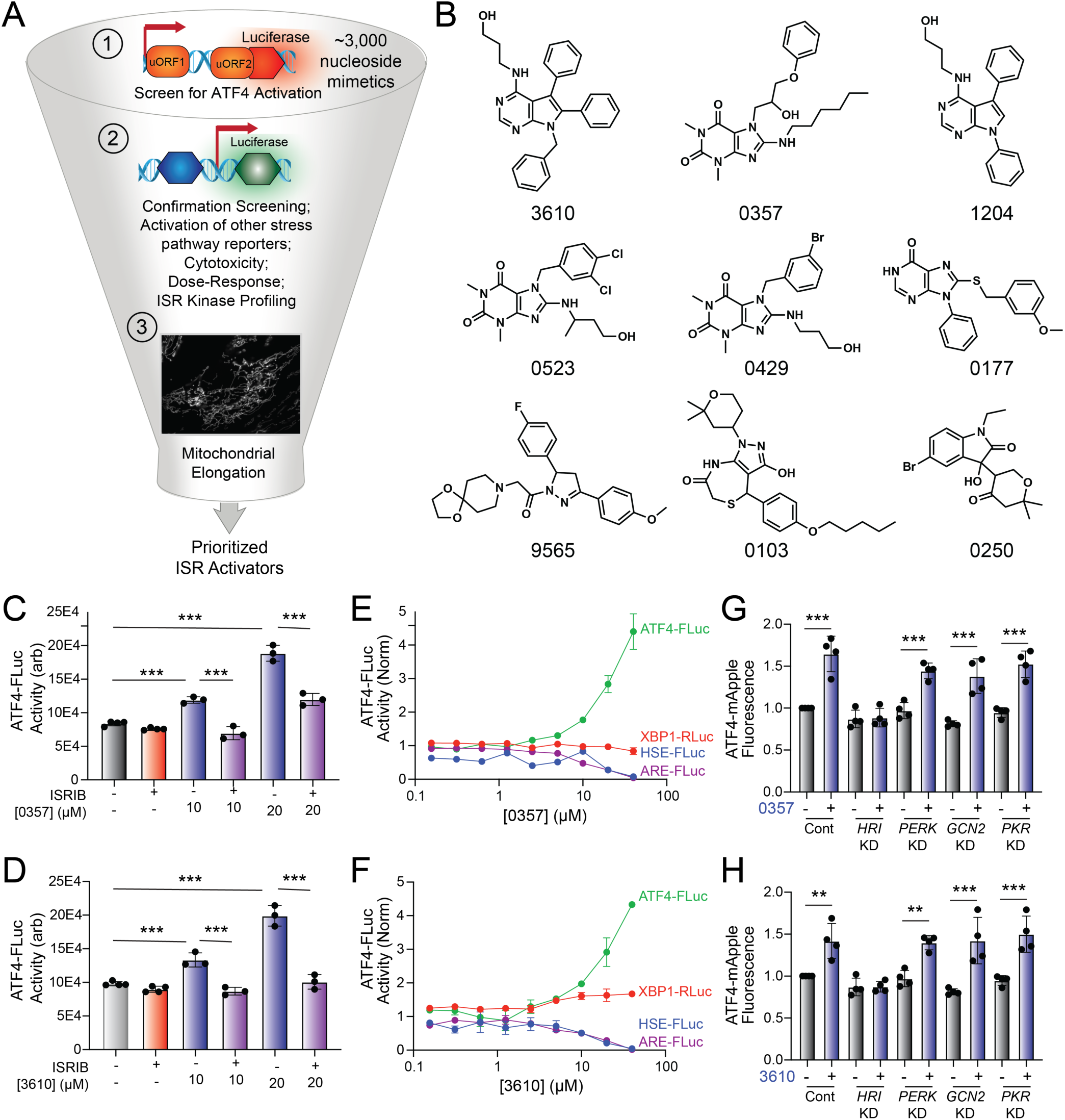
Identification of nucleoside mimetics that prefrentially activate the ISR kinase HRI. **A.** Screening pipeline used to identify selective ISR kinase activating compounds that promote protective mitochondrial elongation. **B**. Structures of the top 9 ISR activating compounds identified in our nucleoside mimetic screen. **C,D**. ATF4-FLuc activity in HEK293 cells stably expressing ATF4-FLuc^10^ treated for 8 h with the indicated concentration of 0357 (**C**) or 3610 (**D**) in the absence or presence of ISRIB (200 nM). **E,F**. Activation of the ATF4-FLuc ISR translational reporter (green), the XBP1-RLuc UPR reporter (red), the HSE-FLuc HSR reporter (blue) or the ARE-FLuc OSR reporter (purple) stably expressed in HEK293 cells treated with the indicated concentration of 0357 (**E**) or 3610 (**F**) for 16 h. **G,H.** ATF4-mAPPLE fluorescence in HEK293 cells stably expressing ATF4-mAPPLE and CRISPRi depleted of the indicated ISR kinase^8^ treated for 8 h with 0357 (**G**, 20 µM) or 3610 (**H**, 20 µM). **p<0.01, ***p<0.005 for one-way ANOVA.

Next, we sought to identify the specific ISR kinases responsible for ISR activation induced by these two nucleoside mimetics. We monitored the compound-dependent activation of an ATF4-mAPPLE fluorescent reporter stably expressed in HEK293 cells CRISPRi-depleted of each individual ISR kinase (**Fig. S1A**).^8,23^ We previously used this assay to confirm that halofuginone activates the ISR downstream of GCN2 and BtdCPU activates the ISR downstream of HRI.^23^ Treatment with either 0357 or 3610 activates the ATF4-mAPPLE reporter in control cells (**Fig. 1G,H**). CRISPRi-depletion of *HRI*, but no other ISR kinase, blocked ATF4-FLuc activation induced by these two compounds. This finding indicates that these compounds activate the ISR downstream of HRI. Collectively, these results identify 0357 and 3610 as nucleoside mimetic compounds that preferentially activate the ISR through a mechanism involving the ISR kinase HRI.

### Pharmacologic HRI activators promote mitochondrial elongation

Halofuginone-dependent activation of GCN2 promotes adaptive mitochondrial elongation.^23^ However, it is currently unclear if pharmacologic activation of other ISR kinases can similarly induce mitochondrial elongation. Here, we tested the ability of our HRI-activating compounds 0357 and 3610 to induce mitochondrial elongation downstream of the ISR. Previously, ISR-dependent mitochondrial elongation was quantified by manually classifying cells as containing fragmented, tubular, or elongated networks.^19,20^ However, since 0357 and 3610 activate ISR signaling to lower levels than that observed for other compounds (e.g., halofuginone), we posited that these compounds may induce more modest mitochondrial elongation that may be difficult to quantify using this manual approach. To address this, we implemented an automated image analysis pipeline using Imaris software to quantify mitochondrial elongation in MEF cells stably expressing mitochondrial-targeted GFP (^mt^GFP) treated with our compounds (**Fig. S2A**).^57^ We collected Z-stack confocal images of ^mt^GFP-expressing MEF cells (MEF^mtGFP^) and processed images using a deconvolution filter in FIJI to reduce the background and enhance the fluorescent signal. We used the “Surfaces” module on Imaris to generate three-dimensional (3D) segmentation models of mitochondria visible in the deconvolved Z-stacks. Using the surfaces module, we quantified parameters defining mitochondrial shape, including bounding box length, sphericity, and ellipsoid principal axis length (**Fig. S2A**). Treatment with conditions that induce mitochondrial elongation, such as the ER stressor thapsigargin (Tg) and the GCN2 activator halofuginone (HF), increased bounding box and ellipsoid principal axis length, while reducing sphericity (**Fig. 2A-D**, **Fig. S2B-D**) – all changes consistent with increases in mitochondrial length. In contrast, treatment with conditions that promote mitochondrial fragmentation, such as the mitochondrial uncouplers BtdCPU and CCCP (both compounds that activate HRI)^6,23^, reduced bounding box length and ellipsoid principal axis length, while increasing sphericity (**Fig. 2A-D**, **Fig. S2B-D**) – all changes consistent with increased mitochondrial fragmentation.

**Figure 2.**
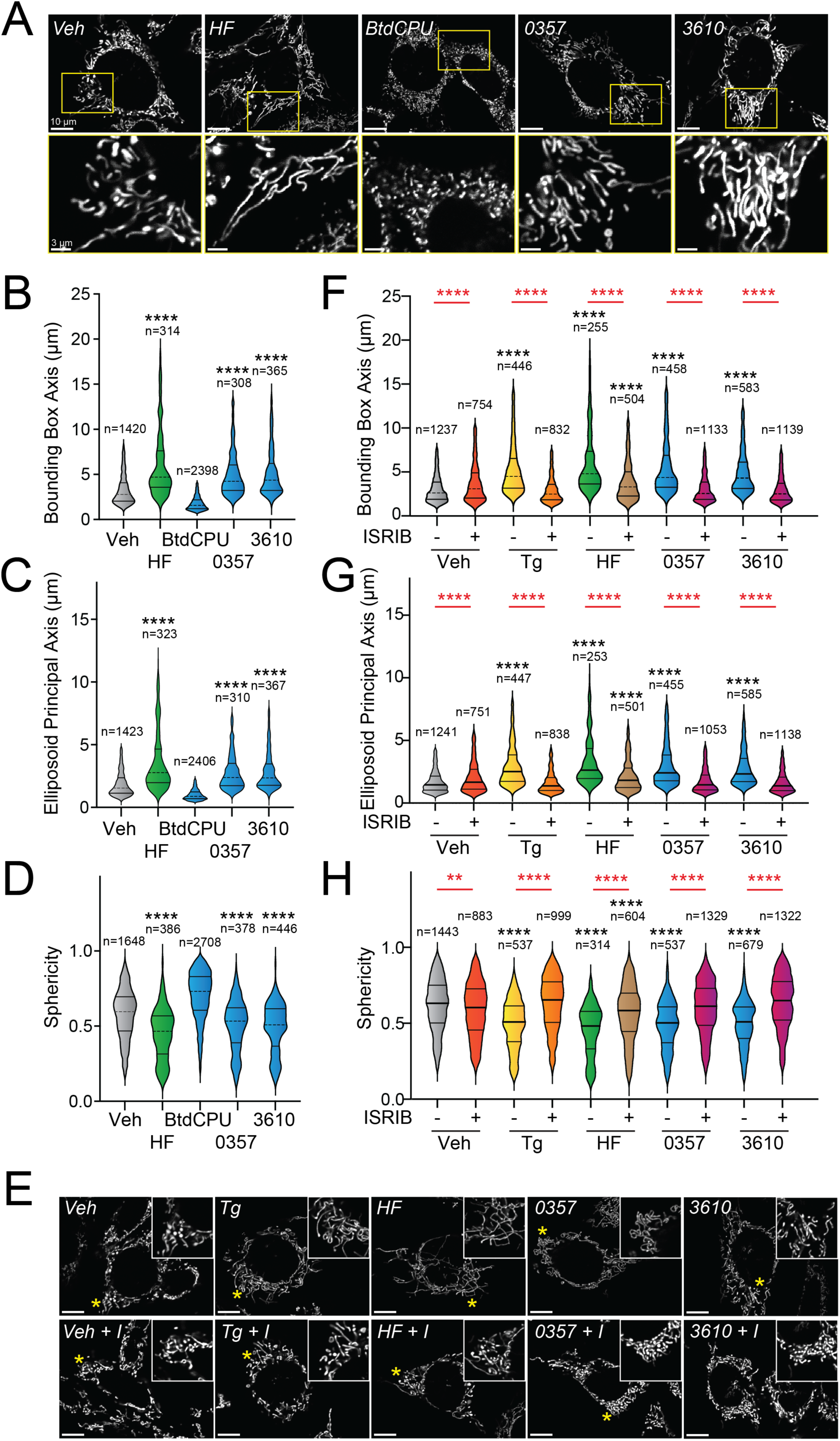
Pharmacologic HRI activation induces ISR-dependent mitochondrial elongation. **A.** Representative images of MEF cells stably expressing ^mt^GFP (MEF^mtGFP^)^57^ treated for 6 h with vehicle (veh), halofuginone (HF, 100 nM), BtdCPU (10 µM), 0357 (10 µM) or 3610 (10 µM). The inset shows a 3-fold magnification of the region indicated by the yellow box. Scale bars, 10 µm (top) and 3.33 µm (bottom) **B-D.** Quantification of bounding box axis, ellipsoid principal axis, and sphericity from the entire dataset of representative images shown in (**A**). The number of individual measurements for each condition are shown above. **E.** Representative images of MEF^mtGFP^ cells treated for 6 h with vehicle (veh), thapsigargin (Tg; 0.5 µM) halofuginone (HF, 100 nM), 0357 (10 µM) or 3610 (10 µM) in the presence or absence of ISRIB (200 nM). The inset shows 3-fold magnification of the image centered on the asterisks. Scale bars, 10 µm. **F-H.** Quantification of bounding box axis, ellipsoid principal axis, and sphericity from the entire dataset of representative images shown in (**E**). The number of 3D segmentations used for the individual measurements for each condition are shown above. *p<0.05, ****p<0.001 for Kruskal-Wallis ANOVA. Black asterisks indicate comparison to vehicle treated cells. Red asterisks show comparisons for ISRIB co-treatment.

We next applied this approach to define the impact of our pharmacologic HRI activators on mitochondrial morphology. Treatment with either 0357 or 3610 for 6 h increased both the bounding box and ellipsoid principal axis length, while decreasing mitochondrial sphericity (**Fig. 2A-D**). This result indicates that both these compounds induced mitochondrial elongation. Co-treatment with the selective ISR inhibitor ISRIB blocked these changes in mitochondria shape, indicating that these compounds induce mitochondrial elongation through an ISR-dependent mechanism (**Fig. 2E-H**). ISRIB co-treatment also blocked mitochondrial elongation induced by the ER stressor thapsigargin (Tg) and the GCN2 activator halofuginone (HF), as predicted.^19,23^ These results indicate that, like halofuginone, pharmacologic HRI activators also induce adaptive, ISR-dependent mitochondrial elongation. Further, these results suggest that the pharmacologic activation of different ISR kinases can induce protective elongation of mitochondria in the absence of cellular stress.

### Pharmacologic ISR activation suppresses ionomycin-induced mitochondrial fragmentation

The ability for our pharmacologic GCN2 or HRI activators to induce adaptive mitochondrial elongation suggests that enhancing signaling through these kinases may suppress mitochondrial fragmentation induced by pathologic insults such as calcium dysregulation.^58–61^ Treatment with the calcium ionophore, ionomycin, induces rapid, DRP1-dependent mitochondrial fragmentation in cell culture models.^46,47^ We pre-treated MEF^mtGFP^ cells for 6 h with the GCN2 activator halofuginone or our two HRI activating compounds (0357 and 3610) and subsequently challenged these cells with ionomycin. We then monitored mitochondrial morphology over a 15-minute timecourse. As expected, ionomycin rapidly increased the accumulation of fragmented mitochondria in these cells, evidenced by reductions in both bounding box and ellipsoid principal axis length and increases of organelle sphericity (**Fig. S3A-D**). Pre-treatment with the ER stressor thapsigargin, which promotes stress-induced mitochondrial elongation downstream of the PERK ISR kinase, reduced the accumulation of fragmented mitochondria in ionomycin-treated cells (**Fig. 3A-D**), as previously reported.^20^ Intriguingly, treatment with halofuginone, 0357, or 3610 also reduced the accumulation of fragmented mitochondria in ionomycin-treated cells. Instead, mitochondria in cells pretreated with these ISR kinase activators and challenged with ionomycin demonstrated mitochondrial lengths and sphericity similar to that observed in vehicle-treated MEF^mtGFP^ cells (**Fig. 3A-D**). These results indicate that pharmacologic activation of different ISR kinases can suppress the accumulation of fragmented mitochondria following ionomycin-induced calcium dysregulation.

**Figure 3.**
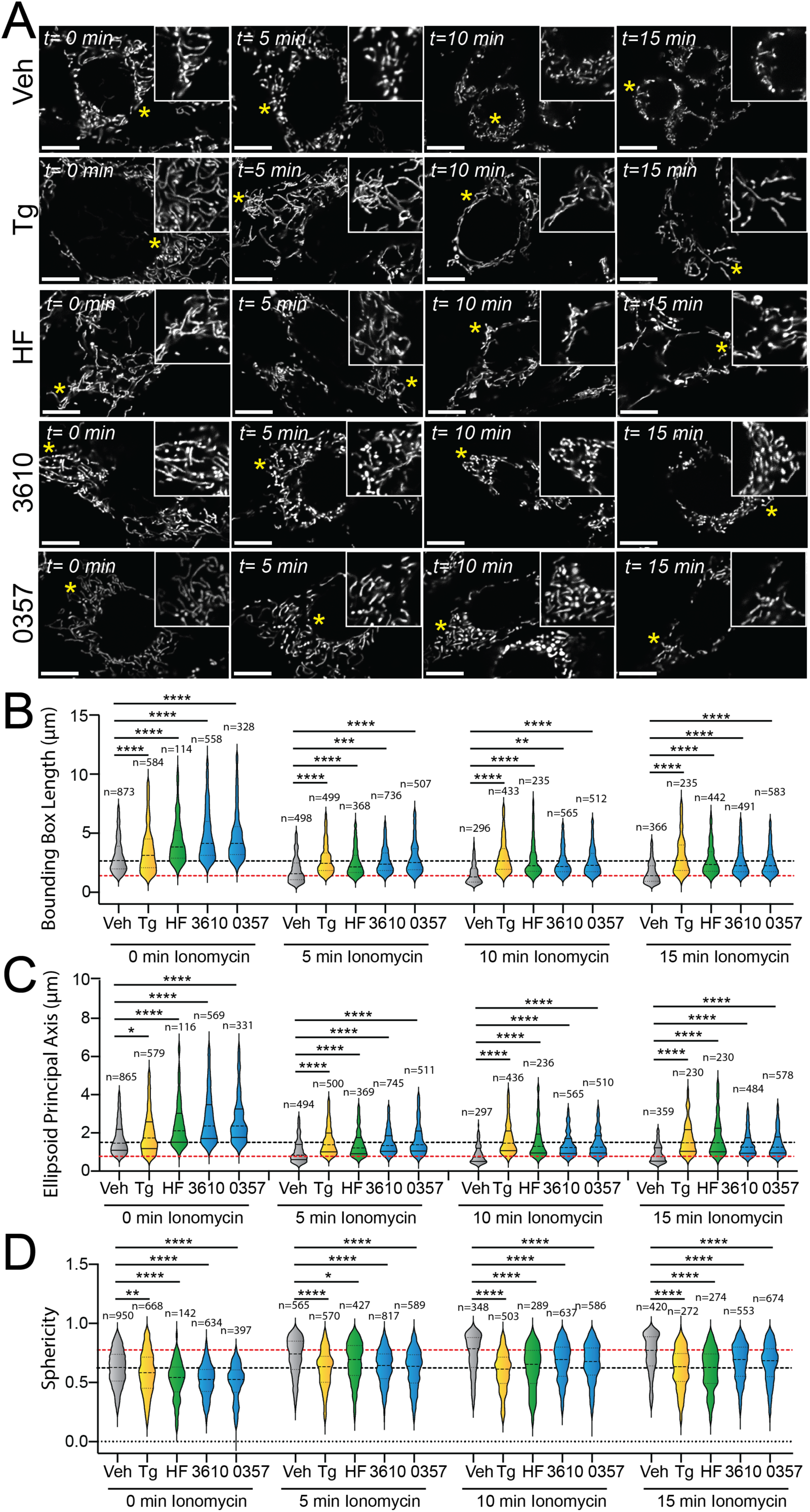
Pharmacologic activation of ISR kinases prevents ionomycin-dependent accumulation of fragmented mitochondria. **A.** Representative images of MEF^mtGFP^ cells pre-treated for 6 h with vehicle (veh), thapsigargin (Tg, 500 nM), halofuginone (HF, 100 nM), 0357 (10 µM) or 3610 (10 µM) and then challenged with ionomycin (1 µM) for the indicated time. The inset shows 3-fold magnification of the image centered on the asterisk. Scale bars, 10 µm. **B-D.** Quantification of bounding box axis length, ellipsoid principal axis length, and sphericity from the entire dataset of representative images shown in (**A**). The black dashed line shows the mean value of vehicle-treated cells prior to ionomycin treatment. The dashed red line shows the mean value of vehicle-treated cells following 15 min treatment with ionomycin. The number of 3D segmentations used for the individual measurements for each condition are shown above. *p<0.05, **p<0.01, ***p<0.005, ****p<0.001 for Kruskal-Wallis ANOVA. Black asterisks show comparison with vehicle-treated cells.

### Pharmacologic activation of ISR kinases restores basal mitochondrial morphology in patient fibroblasts expressing disease-associated MFN2^D414V^

Over 150 pathogenic variants in the pro-fusion GTPase MFN2 are causatively associated with the autosomal dominant peripheral neuropathy Charcot-Marie-Tooth Type 2A.^62–64^ While these pathogenic variants can impact diverse aspects of mitochondrial biology^63^, many, including D414V, lead to increases in mitochondrial fragmentation.^48,62–64^ This can be attributed to reduced activity of MFN2-dependent fusion associated with these variants and a subsequent relative increase of DRP1-dependent mitochondrial fission. We predicted that pharmacologic activation of ISR kinases could rescue mitochondrial network morphology in patient fibroblasts expressing the disease-associated MFN2 variant D414V (MFN2^D414V^). To test this, we treated wild-type human fibroblasts and patient fibroblasts expressing MFN2^D414V^ with halofuginone or our two HRI activating compounds 0357 and 3610 and monitored mitochondrial network morphology by staining with MitoTracker. As reported previously, MFN2^D414V^-expressing fibroblasts showed shorter, more fragmented mitochondrial networks, as compared to control fibroblasts, reflected by reductions in both bounding box and ellipsoid principle axis lengths and increased sphericity (**Fig. 4A,E, S4A-C**).^48^ Treatment with halofuginone, 0357, or 3610 increased mitochondrial length in control fibroblasts (**Fig. 4A-D**). These changes were inhibited by co-treatment with ISRIB, confirming these effects can be attributed to ISR activation. Intriguingly, all three compounds also increased mitochondrial length and reduced sphericity in MFN2^D414V^-expressing patient fibroblasts to levels similar to those observed in control fibroblasts, with halofuginone showing the largest effect (**Fig. 4E-H**). Again, this increase in mitochondrial elongation was reversed by co-treatment with ISRIB. These results show that pharmacologic activation of different ISR kinases can rescue basal mitochondrial morphology in patient fibroblasts expressing the disease-associated MFN2^D414V^ variant that causes dysregulation in various neurological functions including ataxia, optic atrophy, and sensorineural hearing loss.

**Figure 4.**
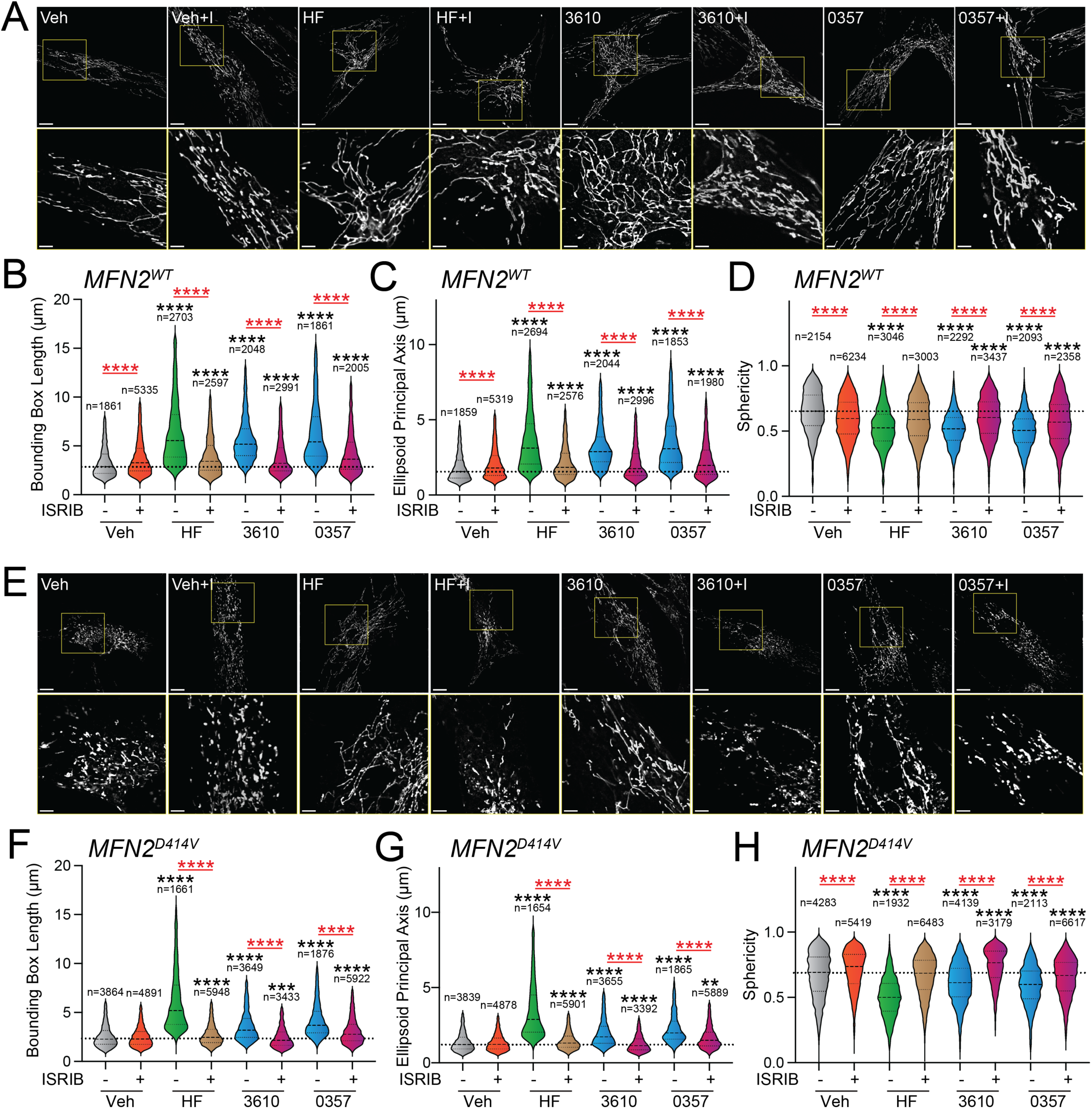
Pharmacologic activation of ISR kinases rescues basal mitochondrial morphology in patient fibroblasts expressing the disease-associated D414V MFN2 variant. **A.** Representative images of control human fibroblasts expressing MFN2^WT^ treated for 6 h with vehicle (veh), halofuginone (HF, 100 nM), 3610 (10 µM), 0357 (10 µM), and/or ISRIB (200 nM). The inset shows 3-fold magnification of the image centered on the asterisks. Scale bars, 15 µm (top) and 5 µM (bottom). **B-D**. Quantification of bounding box axis (**B**), ellipsoid principal axis (**C**), and sphericity (**D**) from the entire dataset of images described in panel **A**. The number of individual measurements for each condition are shown above. **E.** Representative images of patient fibroblasts expressing MFN2^D414V^ treated for 6 h with vehicle (veh), halofuginone (HF, 100 nM), 3610 (10 µM), 0357 (10 µM), and/or ISRIB (200 nM). The inset shows 3-fold magnification of the image centered on the asterisks. Scale bars, 15 µm (top) and 5 µM (bottom). **F-H.** Quantification of bounding box axis (**F**), ellipsoid principal axis (**G**), and sphericity (**H**) from the entire dataset of images described in panel **E**. The number of 3D segmentations used for the individual measurements for each condition are shown above. *p<0.05, ***p<0.005, ****p<0.001 for Kruskal-Wallis ANOVA. Black asterisks show comparison with vehicle-treated cells. Red asterisks show comparisons for ISRIB co-treatment.

## DISCUSSION

Here, we show that pharmacologic activation of different ISR kinases can prevent mitochondrial fragmentation induced by disease-relevant chemical or genetic insults. We identify two nucleoside mimetic compounds that preferentially activate the ISR downstream of the ISR kinase HRI. We demonstrate that these two HRI activators promote adaptive, ISR-dependent mitochondrial elongation. Further, we show that treatment with these HRI-activating compounds or the GCN2 activator halofuginone prevents the accumulation of fragmented mitochondria induced by Ca^2+^ dysregulation and rescues mitochondrial network morphology in patient fibroblasts expressing the disease-associated, pathogenic D414V variant of MFN2. Collectively, these results demonstrate that pharmacologic activation of different ISR kinases can mitigate pathologic mitochondrial fragmentation induced by diverse disease-associated pathologic insults.

Mitochondrial fragmentation is often induced through a mechanism involving stress-dependent increases in the activity of the pro-fission GTPase DRP1.^21,28^ This has motivated efforts to identify pharmacologic inhibitors of DRP1 to prevent mitochondrial fragmentation and subsequent organelle dysfunction associated with human disease.^31,38,65^ Activation of the ISR kinase PERK during ER stress promotes mitochondrial elongation through a mechanism involving inhibition of DRP1.^20^ This suggested that pharmacologic activation of other ISR kinases could also potentially inhibit DRP1 and block pathologic mitochondrial dysfunction induced by increased DRP1 activity. Consistent with this idea, we show that pharmacologic activation of the ISR using both GCN2 and HRI activating compounds blocks the accumulation of fragmented mitochondria in ionomycin-treated cells – a condition that promotes mitochondrial fragmentation through increased DRP1-dependent fission.^46,47^ These results support a model whereby pharmacologic activation of different ISR kinases promotes organelle elongation by inhibiting DRP1 and suggests that pharmacologic activation of ISR kinases can be broadly applied to quell mitochondrial fragmentation in the myriad diseases associated with overactive DRP1.

Pathogenic variants in mitochondrial-targeted proteins are causatively associated with the onset and pathogenesis of numerous neurodegenerative disorders, including Charcot-Marie-Tooth (CMT), spinocerebellar ataxia (SCA), and spastic paraplegia.^21,24,66–69^ In many of these diseases, mitochondrial fragmentation is linked to pathologic mitochondrial dysfunctions, such as reduced respiratory chain activity, decreased mtDNA, and increased apoptotic signaling. Intriguingly, pharmacologic or genetic inhibition of mitochondrial fragmentation mitigates cellular and mitochondrial pathologies associated with many of these diseases, highlighting the critical link between mitochondrial fragmentation and disease pathogenesis.^33–35,43^ Herein, we show that pharmacologic activation of the ISR mitigates mitochondrial fragmentation in patient fibroblasts homozygous for the disease-associated D414V MFN2 variant. This finding highlights the potential for pharmacologic ISR activation to restore basal mitochondrial network morphology in cells expressing pathogenic MFN2 variants, although further study is necessary to determine the impact of ISR activation in cells from heterozygous patients expressing pathogenic MFN2 variants associated with CMT2A, the most common clinical phenotype associated with this disease. Regardless, our results are of substantial interest because over 70% of cases of axonal CMT are associated with pathogenic MFN2 variants, and to date, no disease-modifying therapies are available for any genetic subtype of CMT.^70,71^ Apart from MFN2 variants, our results also highlight the potential for pharmacologic ISR kinase activation to broadly mitigate pathologic mitochondrial fragmentation associated with the expression of disease-related, pathogenic variants of other mitochondrial proteins, which we are continuing to explore.

Apart from morphology, ISR signaling also promotes adaptive remodeling of many other aspects of mitochondrial biology, including proteostasis, electron transport chain activity, phospholipid synthesis, and apoptotic signaling.^13–20^ While we specifically focus on defining the potential for pharmacologic activation of ISR kinases to rescue mitochondrial morphology in disease models, the potential of ISR activation to promote adaptive remodeling of other mitochondrial functions suggests that pharmacologic activators of ISR kinases could more broadly influence cellular and mitochondrial biology in disease states through the adaptive remodeling of these other pathways. For example, we previously showed that halofuginone-dependent GCN2 activation restored cellular ER stress sensitivity and mitochondrial electron transport chain activity in cells deficient in the alternative ISR kinase PERK^23^ – a model of neurodegenerative diseases associated with reduced PERK activity such as PSP and AD.^72,73^ Compounds that activate other ISR kinases are similarly predicted to promote the direct remodeling of mitochondrial pathways to influence these and other functions. Thus, as we, and others, continue defining the impact of pharmacologic ISR kinase activation on mitochondrial function, we predict to continue revealing new ways in which activation of ISR kinases directly regulates many other aspects of mitochondrial function disrupted in human disease.

Several different compounds have previously been reported to activate specific ISR kinases.^23,74–79^ However, the potential for many of these compounds to mitigate mitochondrial dysfunction in human disease is limited by factors including lack of selectivity for the ISR or a specific ISR kinase, off-target activities, or low therapeutic windows.^23^ For example, the HRI activator BtdCPU activates the ISR through a mechanism involving mitochondrial uncoupling and subsequent mitochondrial fragmentation.^23^ Here, we identify two nucleoside mimetic compounds that activate the ISR downstream of HRI and show selectivity for the ISR relative to other stress-responsive signaling pathways, providing new tools to probe mitochondrial remodeling induced by ISR kinase activation. However, the low potency of these compounds limits their translational potential to mitigate mitochondrial dysfunction in disease, necessitating the identification of new, highly-selective ISR kinase activating compounds. As we and others continue identifying next-generation ISR kinase activators with improved translational potential, it will be important to optimize the pharmacokinetics (PK) and pharmacodynamic (PD) profiles of these compounds to selectively enhance adaptive, protective ISR signaling in disease-relevant tissues, independent of maladaptive ISR signaling often associated with chronic, stress-dependent activation of this pathway.^1,2^ As described previously for activators the IRE1 arm of the UPR, such improvements can be achieved by defining optimized compound properties and dosing regimens to control the timing and extent of pathway activity.^80^ Thus, as we and others continue pursuing pharmacologic ISR kinase activation as a strategy to target mitochondrial dysfunction in disease, we anticipate that we will continue to learn more about the central role for this pathway in adapting mitochondria during stress and establish pharmacologic ISR kinase activation as a viable approach to treat mitochondrial dysfunction associated with etiologically-diverse diseases.

## MATERIALS AND METHODS

### Mammalian Cell Culture

HEK293 cells (purchased from ATCC), HEK293 cells stably expressing XBP1-RLuc ^53,54^, HEK293 cells stably expressing HSE-FLuc, HEK293 cells stably expressing ATF4-mAPPLE (a kind gift from Martin Kampmann’s lab)^10^ and CRISPRi-depleted of individual ISR kinases (*HRI*, *PKR*, *PERK*, *GCN2*; a kind gift from Martin Kampmann’s lab at UCSF)^8^, and MEF^mtGFP^ (a kind gift from Peter Schultz)^57^ were all cultured at 37°C and 5% CO_2_ in DMEM (Corning-Cellgro) supplemented with 10% fetal bovine serum (FBS, Gibco), 2 mM L-glutamine (Gibco), 100 U/mL penicillin, and 100 mg/mL streptomycin (Gibco).

Primary fibroblast cells were isolated from partial thickness skin biopsy, as previously described^48,81^, from a patient who provided written informed consent for research studies using human tissues (University of Calgary Conjoint Research Ethics Board REB17-0850). Cells were cultured in Medium Essential Media (11095080, Gibco), supplemented with 10% Fetal Bovine Serum (12483020, Gibco). Cells were maintained at 37 °C and 5% CO_2_. Clinical information regarding this participant was previously reported and included ataxia, optic atrophy, and sensorineural hearing loss.^48^ Exome sequencing in this participant identified a homozygous c.1241A>T variant in *MFN2* (predicted to cause p.(Asp414Val)) and no other pathogenic variants.

### Compounds and Reagents

Compound used in this study were purchased from the following sources: thapsigargin (Tg; Cat# 50-464-294 Fisher Scientific), ISRIB (Cat # SML0843, Sigma), CCCP (Cat #C2759,Sigma), BtdCPU (Cat #32-489-210MG, Fisher), halofuginone (Cat #50-576-3001, Sigma), and oligomycin A (S1478, Selleck). The nucleoside mimetic library was purchased from Chem Div. Hit compounds were repurchased from Chem Div.

### Measurements of ISR activation in ATF4-reporter cell lines

HEK293 cells stably expressing the ATF4-FLuc, HSE-Fluc, or the ARE-Fluc reporter were seeded at a density of 15,000 cells per well in 384-well white plates with clear bottoms (Greiner). The following day, cells were treated with the indicated compound in triplicate at the indicated concentration for 8h. After treatment, an equal volume of Promega Bright-Glo substrate (Promega) was added to the wells and allowed to incubate at room temperature for 10 minutes. Luminescence was then measured using an Infinite F200 PRO plate reader (Tecan) with an integration time of 1000 ms. This assay was used to both screen the nucleoside mimetic library in triplicate and monitor the activity of hit compounds. HEK293 cells expressing the XBP1-RLuc reporter were tested using an analogous approach to that described above, monitoring RLuc activity using Renilla-Glo reagent (Promega), as previously described.^53^

HEK293 cells stably expressing the ATF4-mApple reporter and CRISPRi-depleted of specific ISR kinases were seeded at a density of 300,000 cells per well in 6-well TC-treated flat bottom plates (Genesee Scientific). Cells were treated the next day for 16 h with compound at the indicated concentration. Cells were then washed twice with phosphate-buffered saline (PBS) and dissociated using TrypLE Express (ThermoFisher). Cells were then resuspended in PBS and 5% FBS to neutralize the enzymatic reaction. Flow cytometry was performed on a Bio-Rad ZE5 Cell Analyzer monitoring mAPPLE fluorescence (568/592 nm) using the 561 nm green-yellow laser in combination with the 577/15 filter. Analysis was performed using FlowJo^TM^ Software (BD Biosciences).

### Quantitative Polymerase Chain Reaction (qPCR)

The relative mRNA expression of target genes was measured using quantitative RT-PCR. Cells were treated as indicated and then washed with phosphate-buffered saline (PBS; Gibco). RNA was extracted using Quick-RNA MiniPrepKit (Zymo Research) according to the manufacturers protocol. RNA (500 ng) was then converted to cDNA using the High-Capacity Reverse Transcription Kit (Applied Biosystems). qPCR reactions were prepared using Power SYBR Green PCR Master Mix (Applied Biosystems), and primers (below) were obtained from Integrated DNA Technologies. Amplification reactions were run in an ABI 7900HT Fast Real Time PCR machine with an initial melting period of 95 °C for 5 min and then 45 cycles of 10 s at 95 °C, 30 s at 60 °C.

qPCR Primers

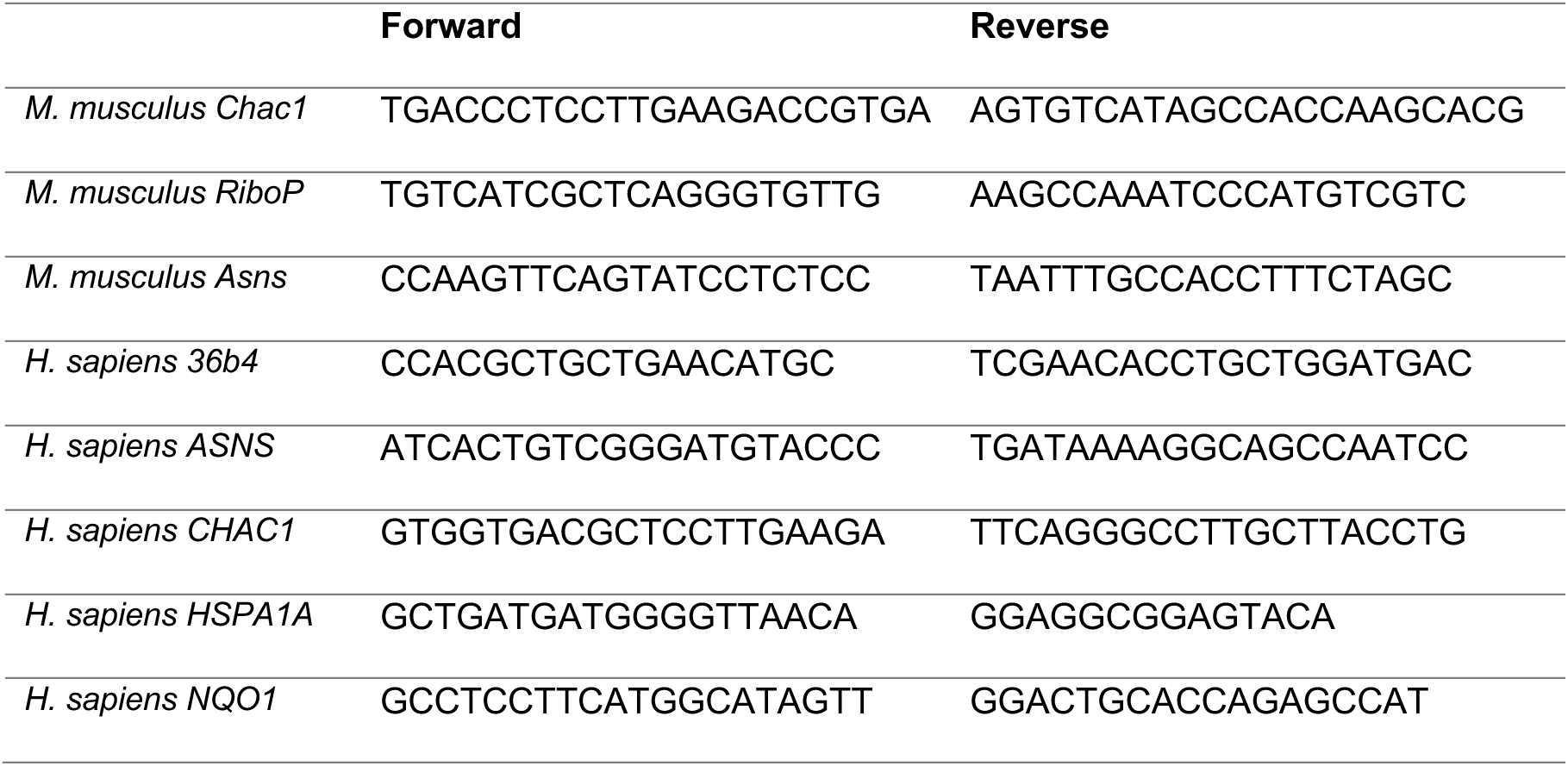

### Fluorescence Microscopy

MEF^mtGFP^ were seeded at a density of 15,000 cells/well in 8-chamber slides (Ibidi) coated with poly-D-lysine (Sigma).^57^ The next day cells were treated with the indicated dose of compound for the indicated time. After treatment, cells were imaged on a Zeiss LSM 880 Confocal Laser Scanning Microscope equipped with a full incubation chamber for regulating temperature and CO_2_ during live cell imaging.

Patient fibroblasts were seeded at 70,000 cells per dish in 35 mm dishes with 20mm glass bottoms (D35-20-1.5-N, Cellvis) for live-cell imaging. After 24 hours, the compound treatments were administered at the dosage and for the time points indicated in the figure legends. The mitochondrial network in patient fibroblasts was visualized using 100 nM MitoTracker Green (M7514, Life Technologies)^82^ for 45 minutes, following washing three times with culture media, according to the manufacturer’s instructions. The Z-stack images were acquired of patient fibroblasts using an Olympus Spinning Disc Confocal System (Olympus SD OSR) equipped with the Olympus UPlanApo 60XTIRF/1.50 Oil Objective using the CellSense Dimensions software. Acquired Z-stacks were analyzed using AI Machine Learning Segmentation (Imaris), as detailed below.

### Quantification of mitochondrial morphology

The z-stack confocal images were processed in FIJI to reduce the background noise and enhance the fluorescent signal. The processed images are then introduced into the developed quantification pipeline in Imaris imaging software. In this approach, mitochondria are segmented in 3D using the “Surfaces” module with a machine-learning algorithm that has been iteratively trained to detect foreground and background pixels in each z-stack, filling blank holes within segments to generate 3D surfaces. The generated surfaces are filtered to include those above a minimum threshold of 250 voxels. While direct length measurements cannot be obtained through the Imaris surface module, indirect measurements of mitochondrial length are inferred from three separate calculations, including (1) object-oriented bounding box axis, (2) ellipsoid axis length, and (3) object sphericity (see **Fig. S2A**). The object-oriented bounding box axis is calculated by measuring the length of the longest or principal bounding-box length of the smallest object-oriented rectangular box that encloses each 3D segmentation. The ellipsoid axis length is calculated by measuring the length of the longest or principal axis of each 3D segmentation. Sphericity is calculated by dividing the longest axis of each 3D segmentation by the length of the perpendicular axis.

### Statistical Methods

Data are presented as mean ± SEM or as violin plots showing the mean and quartiles for the indicated number of measurements. Outliers were removed from datasets describing bounding box length and principal axis length, as appropriate, using the ROUT outlier test in PRISM 10 (GraphPad, San Diego, CA). Normality of datasets from our imaging studies was tested in PRISM 10 (GraphPad, San Diego, CA) using D’Agostino & Pearson, Anderson-Darling, Shapiro-Wilk, and Kolmogorov-Smirnov tests. Statistics were calculated in PRISM 10 (GraphPad, San Diego, CA) and analyzed by one-way ANOVA with Tukey’s multiple correction test, Kruskal-Wallis or Mann-Whitney tests for data exhibiting a non-normal distribution, as indicated in the accompanying figure legends. Indications of nonsignificant interactions were generally omitted for clarity.

## ACKNOWLEDGEMENTS

We thank Jie Sun, and Sergei Kutseikin for experimental support and Evan Powers for critical reading of the manuscript. We thank Kathy Spencer and Scott Henderson in the TSRI Microscopy Facility for their support on the confocal imaging and analysis described in this project. We would also like to thank Martin Kampmann (UCSF), Xiaoyan Guo (UConn), and Jonathan Lin (Stanford) for experimental resources and advice related to this project. This work was supported by the National Institutes of Health (NIH; NS095892, NS125674 to RLW), the Canadian Institutes of Health Research (TES), an NIH F30 Predoctoral Fellowship (AG081061 to KB), National Science Foundation Predoctoral Fellowships (to SO and RA), and the Hotchkiss Brain Institute International Recruitment Scholarship (MZ).

---

**Figure S1.**
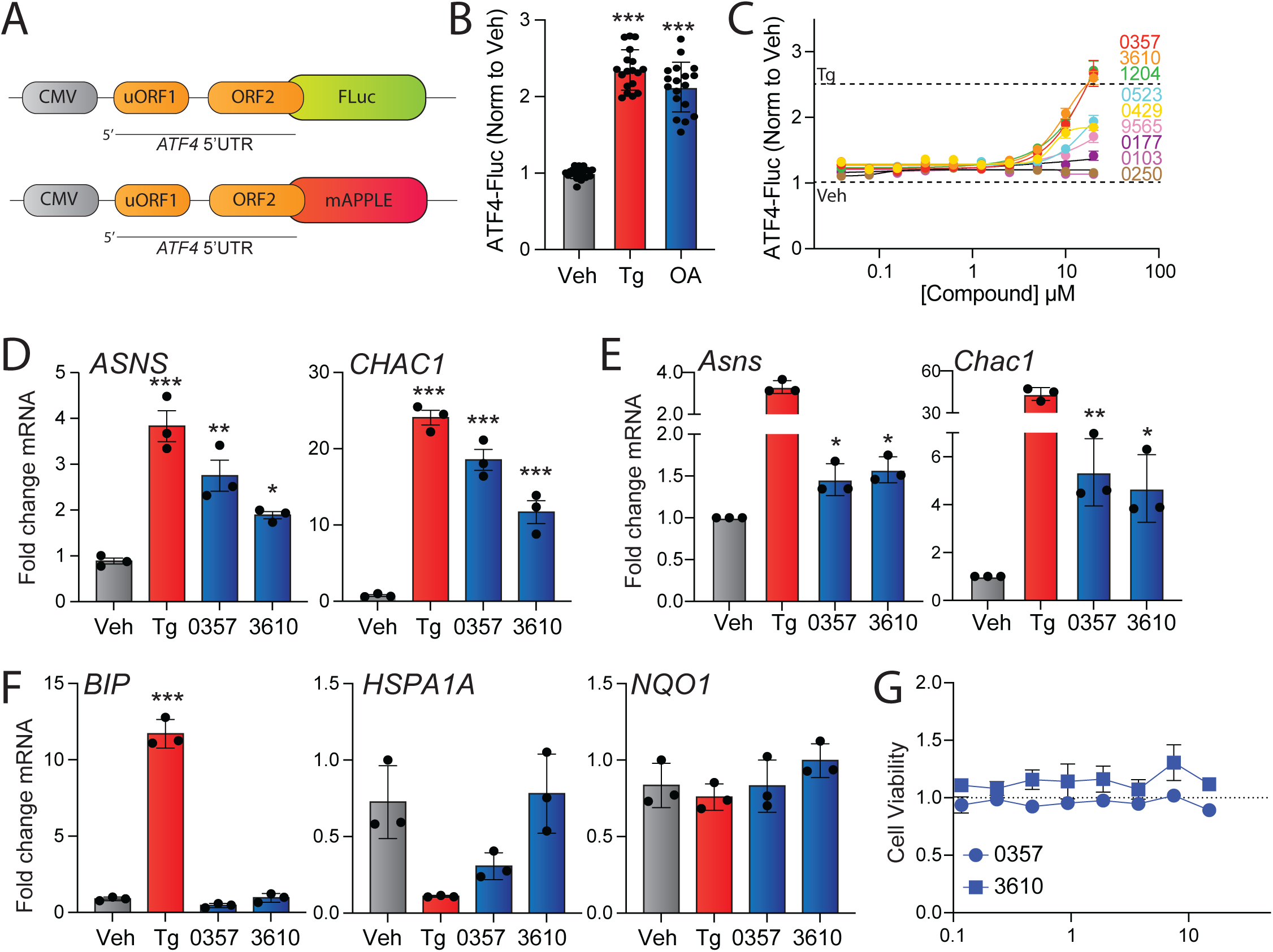
(Supplement to Figure 1). Identification of nucleoside mimetics that preferentially activate the ISR kinase HRI. **A.** The ATF4-FLuc and ATF4-mAPPLE reporters containing the 5’UTR of ATF4.^8,10^ **B**. ATF4-Fluc activity, measured by luminescence, in HEK293 cells stably expressing ATF4-FLuc treated for 8 h with vehicle, thapsigargin (Tg, 0.5 µM), or oligomycin A (OA, 50 ng/mL). **C.** ATF4-FLuc activity, normalized to vehicle, in HEK293 cells stably expressing ATF4-FLuc treated for 8 h with the indicated dose of the indicated compound. The signals observed in veh or thapsigargin (Tg, 0.5 µM) cells is shown by the dashed lines. Error bars show SEM for n=9 replicates. **D**. Expression, measured by qPCR, of the ISR target genes *ASNS* and *CHAC1* in HEK293 cells treated for 8 h with vehicle, thapsgargin (Tg; 0.5 µM), 0357 (25 µM), or 3610 (25 µM). **E**. Expression, measured by qPCR, of the ISR target genes *Asns* and *Chac1* in MEF cells treated for 8 h with vehicle, thapsgargin (Tg; 0.5 µM), 0357 (20 µM), or 3610 (20 µM). **F**. Expression, measured by qPCR, of the UPR target gene *BiP*, the HSR target gene *HSPA1A*, and the OSR target gene *NQO1* in HEK293 cells treated for 8 h with vehicle, thapsgargin (Tg; 0.5 µM), 0357 (25 µM), or 3610 (25 µM). **G.** Viability, measured by Cell Titer Glo, of HEK293 cells treated for 24 h with the indicated concentration of 0357 or 3610. Error bars show SEM for n=3 replicates. *p<0.05, **p<0.01 ***p<0.005 for one-way ANOVA.

**Figure S2.**
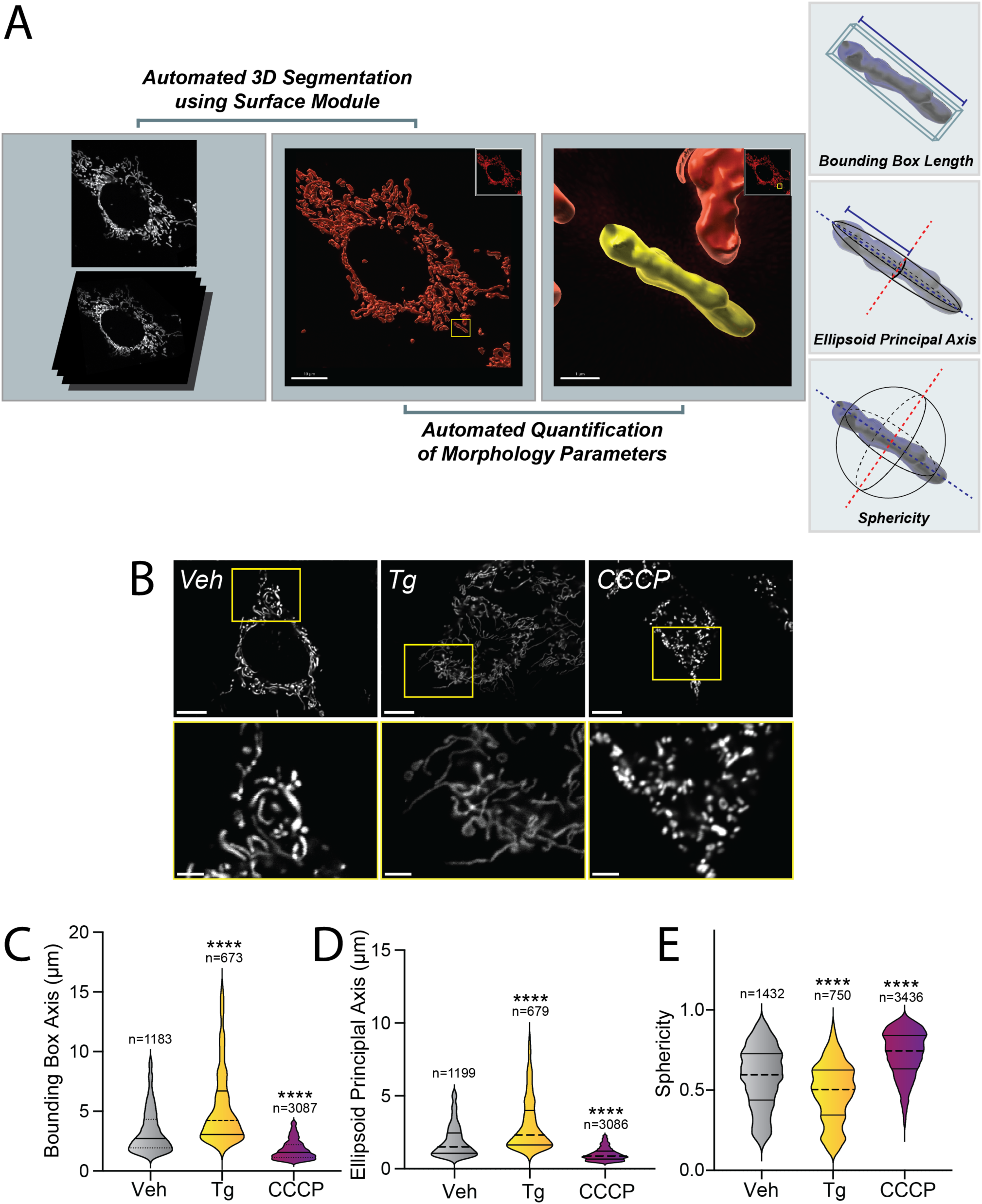
(Supplement Figure 2). Pharmacologic HRI activation induces ISR-dependent mitochondrial elongation. **A.** Image processing and analysis workflow to quantify several parameters that define mitochondrial shape. **B.** Representative images of MEF^mtGFP^ cells treated for 6 h with vehicle (veh), thapsigargin (Tg, 500 nM), or CCCP (10 µM). The inset shows a 3-fold magnification of the region indicated by the yellow box. Scale bars, 10 µm (top) and 3.33 µm (bottom) **C-E.** Quantification of bounding box axis, ellipsoid principal axis, and sphericity from the entire dataset of representative images shown in (**B**). The number of 3D segmentations used for the individual measurements for each condition are shown above. ****p<0.001 for Kruskal-Wallis ANOVA. Black asterisks show comparison with vehicle-treated cells.

**Figure S3.**
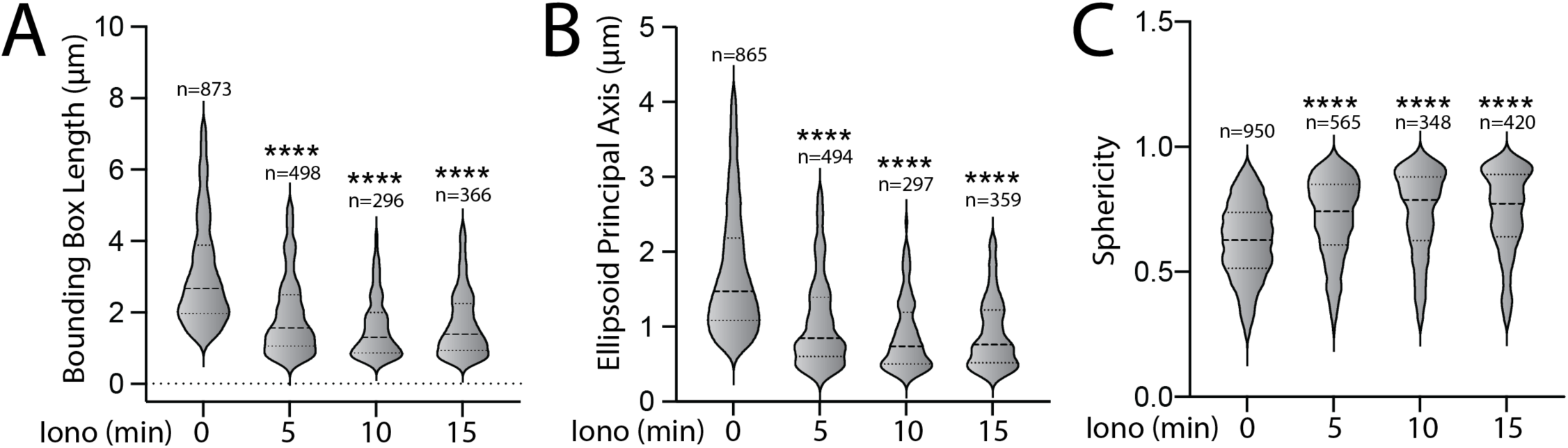
(Supplement to Figure 3). Pharmacologic activation of ISR kinases prevents ionomycin-dependent accumulation of fragmented mitochondria. **A-C.** Quantifications of bounding box axis, ellipsoid principal axis, and sphericity of MEF^mtGFP^ cells pre-treated for 6 h with vehicle and then challenged with ionomycin (1 µM) for the indicated time. Representative images are shown in Fig. 3A. The number of 3D segmentations used for the individual measurements for each condition are shown above. ****p<0.001 for Kruskal-Wallis ANOVA. Black asterisks show comparison with vehicle-treated cells at time 0.

**Figure S4.**
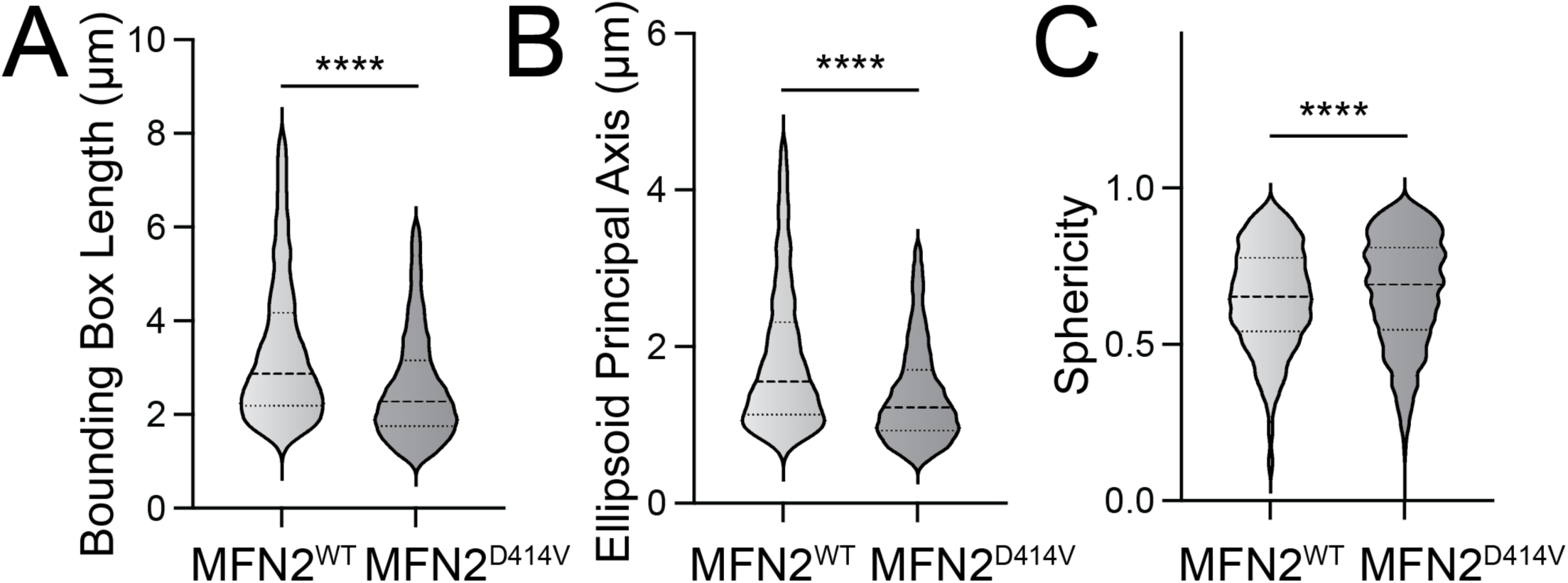
(Supplement to Figure 4). Pharmacologic activation of ISR kinases rescue basal mitochondrial morphology in patient fibroblasts expressing the disease-associated D414V MFN2 variant. **A-C**. Bounding box length (**A**), ellipsoid principal axis length (**B**), and sphericity (**C**) in control human fibroblasts expressing MFN2^WT^ or patient fibroblasts expressing MFN2^D414V^. Representative images are shown in **Fig. 4A,E**. ****p<0.001 for Mann-Whitney t-test.

